# HIV-1 Rev Protein Forms a Zinc-Linked Dimer

**DOI:** 10.1101/2024.12.06.627258

**Authors:** Raza S. Khan, Robert O. Fox

**Affiliations:** Department of Biology & Biochemistry, University of Houston, Houston, TX 77204-5001

## Abstract

HIV-1 Rev protein is a small, basic protein of 116 amino acids that assembles reversibly in the presence and absence of its cognate RRE containing RNAs both *in vivo* and *in vitro*. The biologically active form of Rev is unclear since studies have shown monomer, dimer, tetramer and higher-order oligomer interactions with various RRE-RNAs. Whilst assembly is essential for its regulatory role in the viral life cycle, it has been a barrier to high resolution structural studies of the whole protein and its complexes. The N-terminal half of Rev has been shown to contain a helix loop helix motif with residues involved both in assembly and RNA binding. The C-terminal half is predicted to contain little secondary structure based on UV-CD spectral analyses, and to contain the leucine rich activation domain (residues 73-83). Early studies had shown the essential part of the C-terminal extends to residue 93 and is required for increased structural stability of the protein and its complexes with RRE-RNAs, and to facilitate the formation of Rev dimers (4, 19). The strong conservation of cysteines at positions 85 and 89, and the less well-conserved histidine residues at 53 or 82 led us to examine Rev-metal interactions. Here we show that Rev binds Zn^2+^ with a stoichiometry of one equivalent per Rev dimer. Optical spectroscopy of Rev Co^2+^ complexes revealed that the metal site is composed of four cysteine residues with a tetrahedral coordination geometry. We propose that HIV-1 Rev protein is biologically active as a Zn^2+^Cys4-linked dimer.

---

The HIV-1 Rev protein promotes the synthesis of late gene products by facilitating the nuclear export of singly spliced and unspliced, virally encoded RNAs (*1–3*). Rev binds with high affinity to the Rev Response Element (RRE), a structured region of these RNAs, and assembles on the RNA (*4–6*). This RRE-Rev assembly is critical for Rev function and is reversible, allowing Rev to shuttle between the nucleus and cytoplasm via interactions with cellular importin and exportin (*7*). Rev forms homomultimers *in vivo*, and *in vitro* forms an array of multimers, leading ultimately to formation of filaments at high protein concentrations (*8*). A low-resolution EM structure of Rev filaments identified a Rev dimer as the repeating unit (*9*). Studies of the wild-type protein and defined mutants have shown that Rev can interact with RRE-RNAs as a monomer at low protein concentrations, but stable complexes are formed only when Rev dimerizes on the RRE-RNAs or interacts directly as dimers or higher oligomers (*10, 11*). Genetic, mutational and biochemical studies suggest a model whereby Rev assembles on the RNA via a rotationally symmetrical interaction involving residues in the two N-terminal helices of each monomer (*12*).

Cysteine and histidine side chains are often involved in Zn^2+^ coordination in nucleic acid binding proteins. The conservation of HIV-1 Rev Cys85 and Cys89, and the presence of less conserved histidine residues, lead us to examine Rev for Zn^2+^ binding by flame atomic absorption spectroscopy (AAS) after exhaustive dialysis against various buffers. The wild-type and a dominant-negative mutant RevM10 variant (L78D:E79L) contained 0.51 and 0.56 equivalents of non-dialyzable Zn^2+^, respectively, suggesting a Rev dimer shares a single Zn^2+^ ion under these solution conditions. No significant difference was seen when Zn^2+^ was added before or after the protein had been refolded (0.51 vs 0.61 equivalents respectively), provided the protein was kept under reducing conditions. The bound Zn^2+^ could be removed by dialysis against a buffer containing 1mM EDTA chelator.

Zn^2+^ titrations of RevM10 were monitored by measuring fluorescence quenching of the single tryptophan residue (Trp45). Quenching appeared complete after the addition of 0.7 equivalents of Zn^2+^ (Figure 1). This result also supports the sharing of one Zn^2+^ ion by a Rev dimer. Under these solution conditions Zn^2+^ binding is tight and stoichiometric, indicating Rev affinity for Zn^2+^ is in the submicromolar to micromolar range, and supporting the AAS data above and in Table S1 of the Supporting Information. The extent of fluorescence quenching was modest, and no obvious shift was observed in the tryptophan fluorescence peak, suggesting that there were no major structural changes in the immediate vicinity of the chromophore upon Zn^2+^ binding.

**Figure 1:**
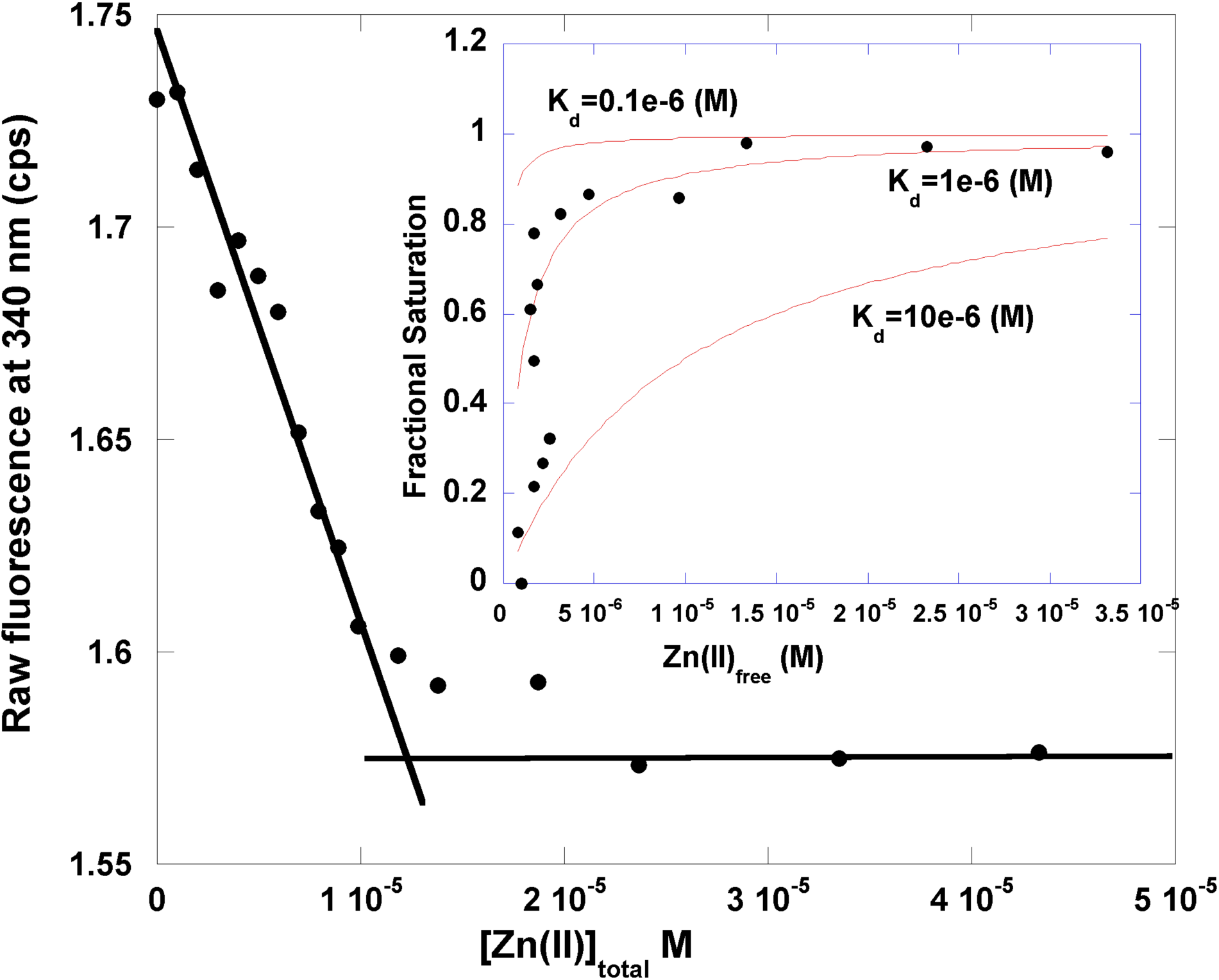
RevM10 binds Zn^2+^. The dominant negative mutant RevM10 displays reduced assembly (Malim et al. 1993) and was used to facilitate Zn^2+^ binding titrations The intrinsic tryptophan fluorescence at 340 nm is plotted added [Zn^2+^]. To 17.3 uM reduced protein in 10 mM Tris, 0.4 M NaCl, pH 7.5, small aliquots of concentrated stock of zinc chloride were added. Excitation was at 275 nm with slit widths set to 0.5 mm for both excitation and emission monochromators. The straight lines drawn through the data suggest a stoichiometry of 0.7 equivalents Zn^2+^ per monomer. Upon continued titration beyond the stoichiometric point light scattering is seen, suggesting increased assembly and linking metal binding to protein assembly. This complicates meaningful analysis and determination of intrinsic metal affinity. However, ignoring the linkage to assembly, the inset shows goodness of three fits to the data over three orders of magnitude.

The identity of protein residues involved in metal coordination, and the geometry of the metal coordination site, can be determined by optical spectroscopy of isomorphous protein-cobalt complexes (*13*). Titration of Rev with Co^2+^ displayed an absorption peak at 307 nm, and a shoulder at 360 nm in the UV region (Figure 2A). These optical features are characteristic of S^-^ ® Co^2+^ ligand to metal charge transfer bands (LMCT), and are considered a signature for thiolate ligands (*13, 14*). In the visible region of the spectrum, absorption bands were seen at ∼620 nm, ∼675 nm and ∼725 nm (Figure 2A and 2C). These features arise from Co^2+^ d-d transitions, and are characteristic of tetrathiolate coordination (*13*). The results indicated that Rev binds Co^2+^ via four Cys residues, excluding involvement of histidines in metal coordination (*15*). The tetrathiolate stoichiometry was confirmed by the magnitude of the extinction coefficient at 307 nm of ∼4000 M^-1^cm^-1^ for the Rev-Co^2+^ complex (Figure 2B). This value of ∼1000 M^-1^cm^-1^ per S^-^→Co^2+^ bond is in the range (900-1300 M^-1^ cm^-1^) found in other thiol-rich proteins (*13, 14*).

**Figure 2:**
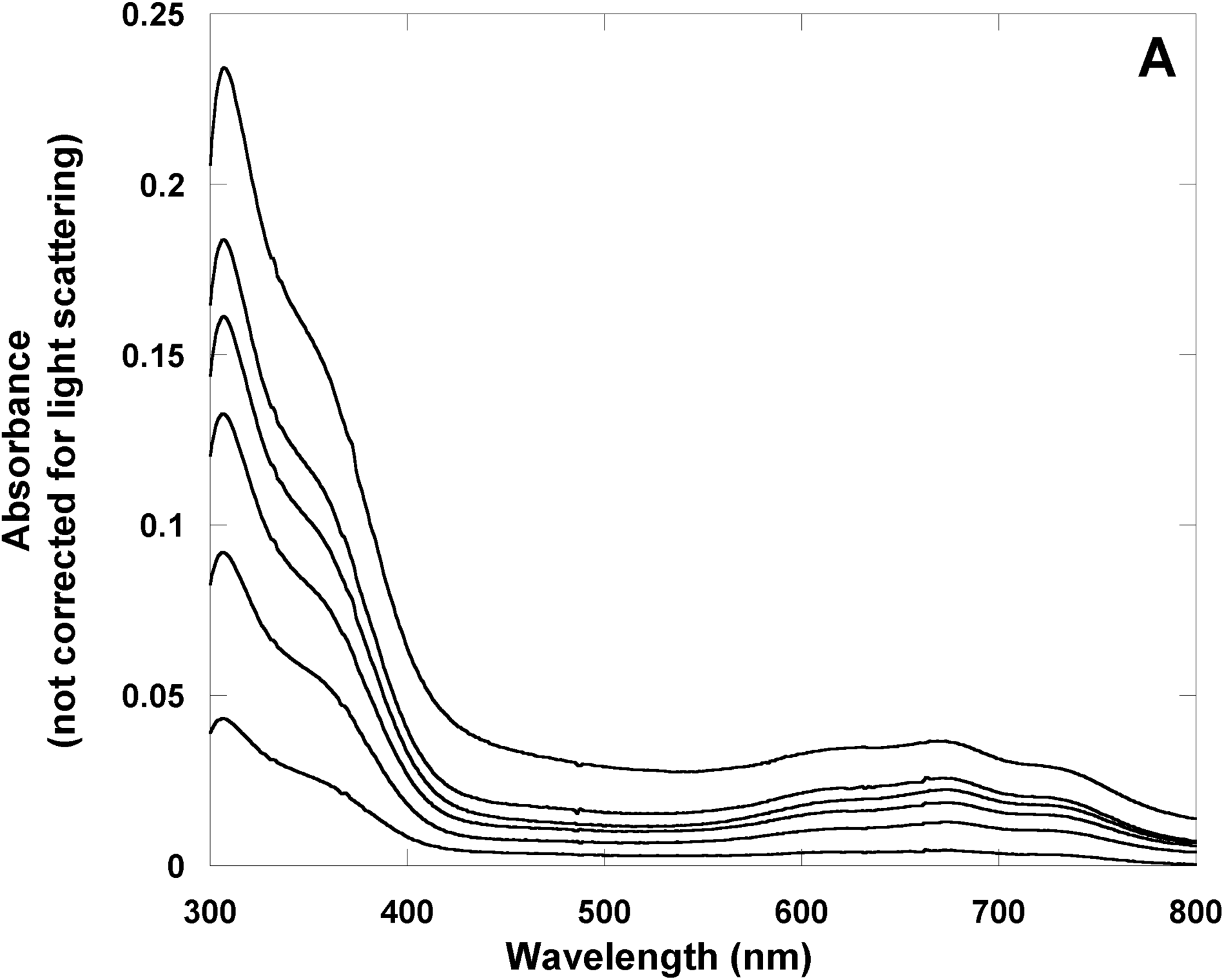
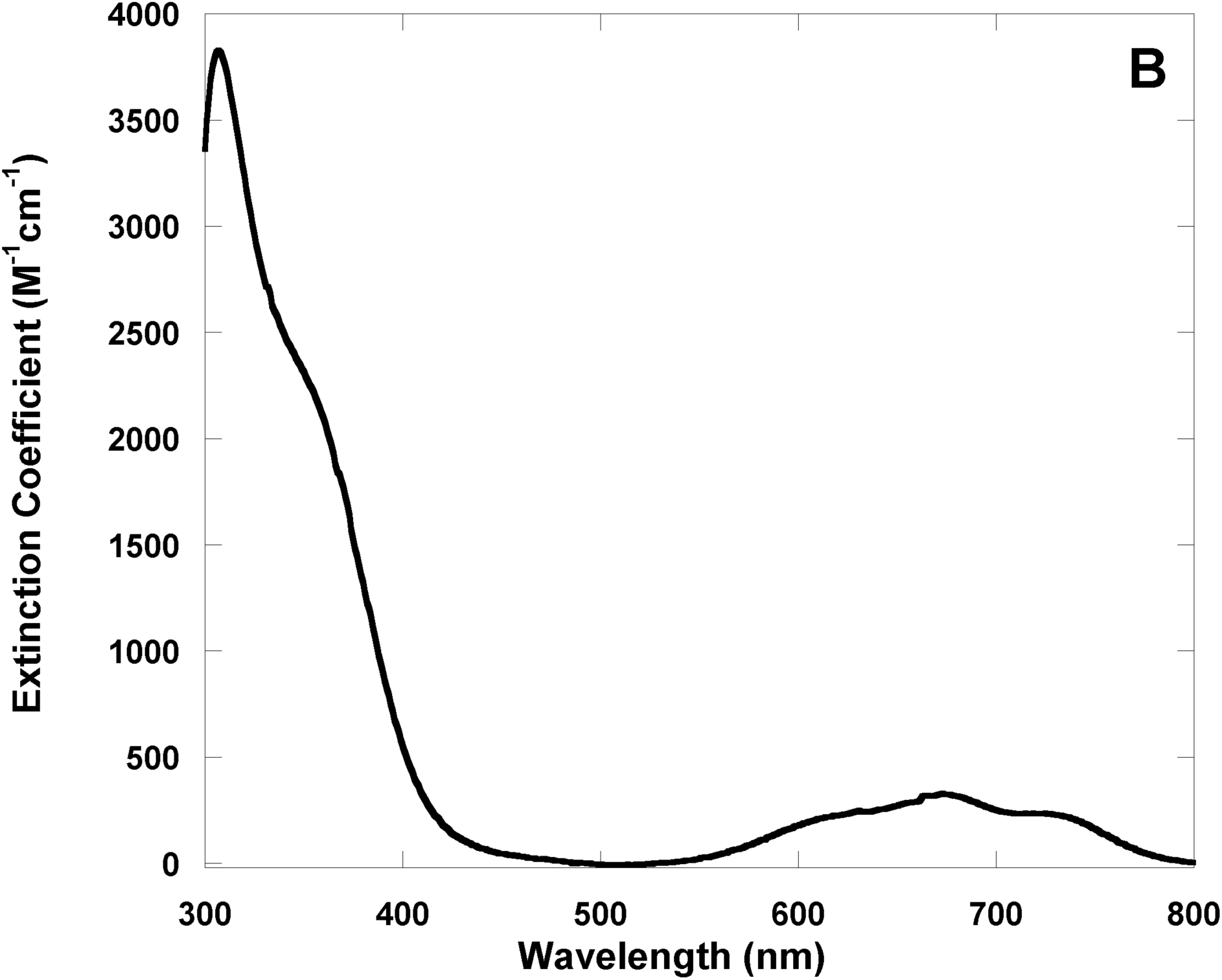
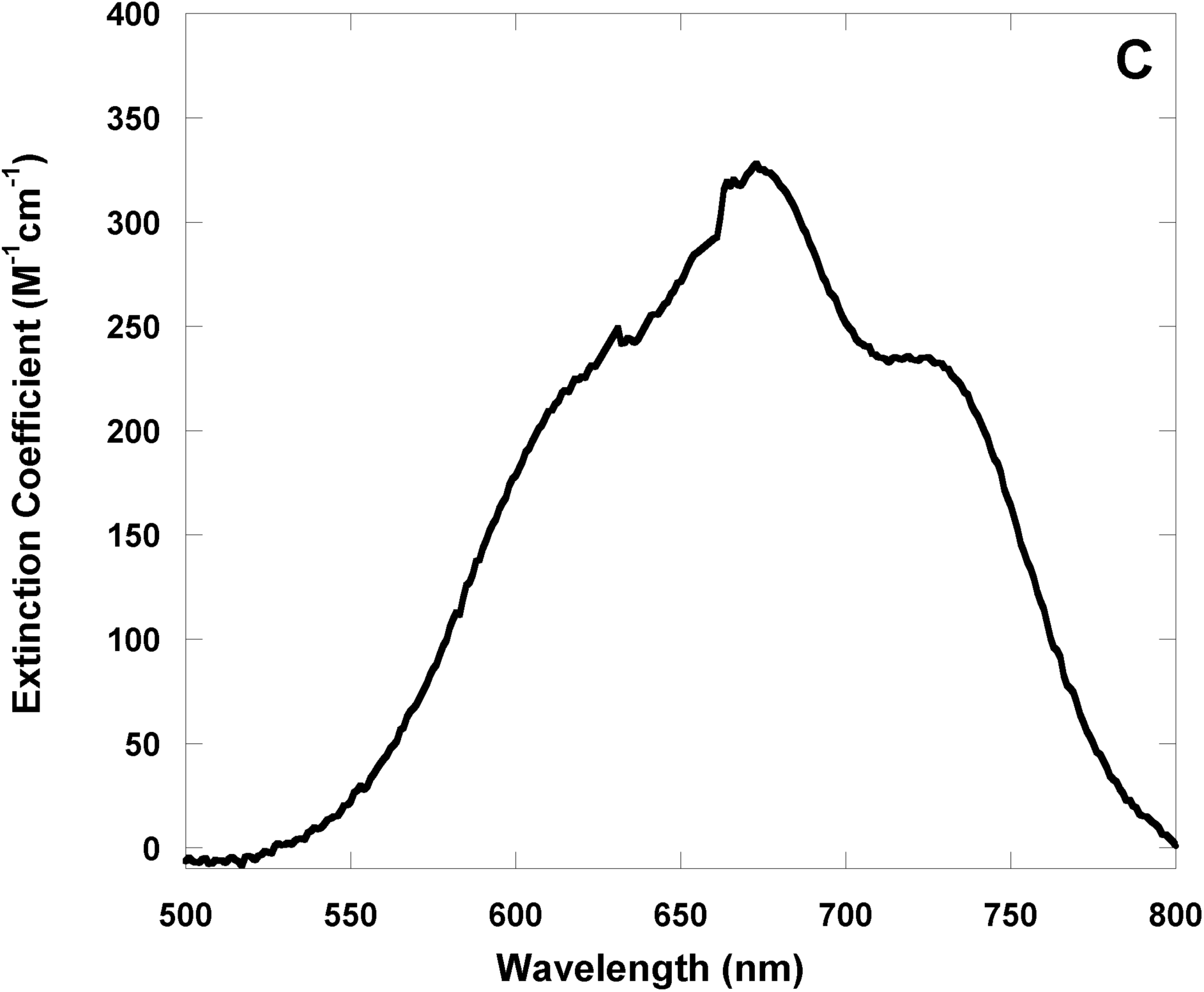
Rev binds Co^2+^. Reduced, unfolded Rev was refolded under anaerobic conditions. 160 uM refolded Rev, (quantitated using Elman’s reagent), in 50mM Tris, 3M NaCl, pH 7.5 was titrated with small aliquots of concentrated Co^2+^ in an anaerobic cuvette. The difference UV-Vis absorbance spectrum at 0.06, 0.12, 0.16, 0.22, 0.29 and 0.47 equivalents of added Co^2+^ is shown in **Panel A**. The arrow indicates increasing [Co^2+^]. The calculated extinction coefficient at 0.29 equivalents of Co^2+^ and corrected for light scattering is plotted against wavelength in the UV region of the absorption spectrum in **Panel B**, and in the visible region in **Panel C**. As the titration proceeds, assembly is promoted as indicated by the apparent increase in absorbance around 500 and 800 nm seen in panel A.

The coordination geometry of tetrathiolate Co^2+^ complexes is revealed by the magnitude of the extinction coefficients of the d-d transition bands in the visible region of the difference absorption spectrum (*13, 14*). Tetrahedral and distorted tetrahedral coordination geometries give extinction coefficient values above 250 M^-1^cm^-1^, while an octahedral geometry would give values of <30 M^-1^cm^-1^ (*13, 14*). The Rev-Co^2+^ complex displayed an extinction coefficient of ∼350 M^-1^cm^-1^ at 675 nm, suggesting a tetrahedral or distorted tetrahedral geometry of this tetrathiolate site (Figure 2C).

The presence of metal should protect the Rev cysteine thiols from air oxidation *in vitro*, and prevent formation of both intra- and inter-molecular disulfides. Rev samples were prepared with and without Zn^2+^, and their thiol content determined over a period of days. Figure S1 shows that in the presence of Zn^2+^, the reduced thiol content remains significantly higher than in the apo-Rev sample (Figure S1 of the Supporting Information). The AAS and the fluorescence quenching titration data demonstrate the stoichiometry is one Zn^2+^ equivalent per Rev dimer. This is confirmed by the optical spectroscopy of Rev-Co^2+^ complex, demonstrating that four thiol ligands are provided by two Rev monomers. Thus, each monomer forms a half-site consisting of Cys85 and Cys89 for metal coordination. At the protein concentrations used for the AAS and Zn^2+^ titrations, Rev exists as dimers and higher order oligomers (*8, 16, 17*). Taken together these data strongly support the existence of Rev as a Zn^2+^Cys_4_ dimer.

Whether the presence of Zn^2+^ is purely structural or is required for one or more Rev functions, is unknown. Given the many interactions of Rev with RRE-containing RNAs and cellular cofactors, and its shuttling activity through various cellular environments, it is possible that Zn^2+^ availability and coordination play a modulating role for some Rev function. Zn^2+^ availability varies during the cell cycle and within cellular compartments; both have been shown to modulate cellular signaling and the activity of Zn^2+^-containing transcription factors (*18*).

Ultracentrifugation studies of purified Rev have shown a distribution of monomers, dimers and higher oligomers that shifts towards higher molecular weight oligomers with increased protein concentrations, and that this assembly is reversible (*8, 16, 17*). It is not clear from these studies if Rev protein assembly proceeds via dimer or monomer oligomerization that leads eventually to the formation of filaments. One low-resolution structure of Rev filament shows the Rev dimer as the unit that assembles (*9*). From various studies it is clear that there is a range of Rev multimers for a given protein concentration, and that the Rev dimer predominates at protein concentrations of about 10 uM (*8, 19*). Our AAS and Zn^2+^-binding experiments were carried out in this concentration range, and a tetrathiolate metal-binding site is formed by Rev dimers. At the higher concentrations used in the Co^2+^ experiments, the tetrathiolate metal-binding site must also be present to bind the metal, even though at these protein concentrations it is expected that high molecular weight oligomers exist. Thus, our results support a model in which Rev dimers assemble into oligomers.

Gel-shift assays and other studies suggest that Rev binds RRE-RNAs as a monomer at low Rev:RNA ratios, but as the ratio increases a ladder of complexes is seen, suggesting that Rev can bind RRE-RNAs as dimer or oligomer (*10, 11, 20, 21*). Solution binding studies also suggest that a second Rev monomer is needed to form a stable, tight-binding complex with RRE-containing RNAs (*11, 20*). It is possible that Zn^2+^ occupancy of this dimerized Rev-RRE complex modulates the interaction and assembly on the RRE.

UV-CD spectra of Zn^2+^Cys_4_ dimer and of the apo-Rev show little difference, suggesting that there was no significant change in the overall secondary structure (data not shown). Since there is also modest quenching of the Trp45 fluorescence and no shift in the emission peak, it seems likely that Zn^2+^ fills a pre-formed metal-binding site. Thus it is likely the cysteines (and possibly the residues flanking them) from each monomer are not only close in space but oriented to form a tetrathiolate and tetrahedral site that Zn^2+^ fills. This would require the cysteine-containing region within the Rev C-terminal half to form a dimer interface, as was previously demonstrated (*19, 20*).

How the activity of the Zn^2+^Cys_4_ dimer differs in one or all of the Rev protein interactions and functions awaits investigation. Rev occupies varied cellular environments, from its synthesis in cytoplasm to assembly and storage in the nucleolus, to its functions of binding RRE-containing RNAs, assembling and transporting these out of the nucleus and into the cytoplasm for translation or packaging, and shuttling back to the nucleolus. The assembled states of the protein vary from monomer to oligomer during this Rev cycle, both when free and bound to RRE-containing RNAs and to cellular cofactors.

Bound Zn^2+^ may facilitate reversible Rev assembly. Since available Zn^2+^ levels are known to vary in the different cellular compartments, and as a function of the cellular state, it is feasible that one or more Rev function is modulated by occupancy of its metal-binding site. Such sensitivity to Zn^2+^ levels could provide a mechanism by which HIV-1 switches to late gene expression that results in virus production.

## ACKNOWLEDGMENT

We thank Drs. T. Daly and J. C. Lee for the Rev proteins. We also thank Dr. D. P. Giedroc for valuable advice and the use of his anaerobic glovebox and AAS, and Dr. D. Konkel for critically reading the manuscript. R.S.K. was supported by a postdoctoral fellowship from the James W. McLauglin Foundation. This work was supported by a Development Grant from the Sealy & Smith Foundation to R.O.F. and a grant from the Welch Foundation (H-1345) to R.O.F.

## SUPPLAMENTARY MATERIALS

### Materials and Methods

#### Rev proteins

Unfolded and partially purified wild-type Rev protein and mutant RevM10 were a gift from Drs. T. Daly and J. Ching Lee. These proteins were purified under denaturing conditions using reverse phase C18 chromatography and refolded on a fast-flow SP-Sepharose column as published, except that Tris-HCl buffers at pH 7.5 were used with the addition of 1 mM EDTA and 2 mM DTT.

Making the Zn^2+^-linked dimer aerobically: As Zn^2+^ does not have multiple oxidation states of Co^2+^, the RevZn^2+^ dimer of the wild type and mutants can be formed aerobically. After the reverse phase chromatography step, Rev proteins are stored lyopholized at −70° C. To refold the Rev proteins small amounts (2-5 mg) of the lyopholized protein are resuspended in a small volume ( up to 0.5 mL) of 50 mM Tris-HCl, 8 M urea and 10 mM DTT pH 7.5 (Buffer A) and incubated at 45° C for about 2 hours. The reduced, unfolded protein is loaded on to a small SP-Sepharose column (∼2-5 mL) previously equilibrated in Buffer A, washed extensively with 50 mM Tris-HCl, 0.4 M NaCl, pH 7.5 and then eluted with 50 mM Tris-HCl, 3 M NaCl, pH 7.5. The eluted protein is quantitated by both absorbance at 280 nm and by Ellman’s reagent and Zn^2+^ is added to 0.6 equivalents, followed by microcentrifugation to remove any precipitation. The protein is quantitated again by absorbance at 280 nm and used. For storage and prolonged use 2 mM DTT is added to the protein stock or dialysis buffers.

#### Fluorescence titrations

Concentrated stocks of purified and refolded proteins were dialyzed and stored in 10 mM Tris-HCl, 2 M NaCl, 2 mM DTT, pH 7.5. For titrations 2.0 mL of protein solutions (2-17 mM) were prepared in 10 mM Tris-HCl, 0.4 M NaCl, pH 7.5. Small aliquots of concentrated Zn^2+^ in water were added with stirring at room temperature, and the fluorescence change detected on a SPEX Fluoromax 2 spectrofluorometer. The same volume of protein dialyzate was diluted to 2.0 mL and titrated with Zn^2+^ and the readings at appropriate Zn^2+^ concentrations were subtracted from the protein fluorescence.

#### Atomic absorption spectroscopy

Various Rev samples were prepared as shown in Table 1 and submitted for Zn^2+^ analyses using a Perkin Elmer Model 5100 spectrometer or carried out on a Perkin Elmer Model 2380 spectrometer operating in the flame mode with detection at 213.7 nm and at a slit setting of 0.7 mm.

#### Thiol determination

Rev was denatured and thiols reduced in the presence of 10 mM Tris-HCl, 8 M urea, 20 mM DTT, pH 7.5, and incubated at 42° C for 2 hours. From this stock, apo-Rev was refolded on a SP-Sepharose column in the absence of Zn^2+^ and DTT using the procedure above. A small aliquot of eluted apo-Rev was quantitated by UV absorbance ((280=8,440 M^-1^cm^-1^). The number of reduced thiols was determined immediately using 5,5í-dithionitrobenzoic acid (DTNB). For thiol determination protein samples (2-10 mM) were diluted into 300 mL of 10 mM Tris-HCl, 1 M NaCl, with or without 8 M urea, pH 7.5, 30 mL of 25 mM DTNB was added and the reaction mix incubated for up to 2 hours. The maximum absorbance at 412 nm was recorded and concentration of thiol determined using ((412=13,600 M^-1^cm^-1^. For Zn^2+^-linked dimer, 100 mM Zn^2+^ was added to the denatured stock, before refolding as above, except all buffers contained 100 mM Zn^2+^. Both forms of Rev were then subjected to exhaustive dialysis into 10 mM Tris, 1 M NaCl, pH 7.5. Thiols were quantitated over a period of 8 days and Zn^2+^ content of both samples was determined by AAS. All values have been subtracted for thiol content of buffers and dialyzates which gave background values of less than 10%.

#### Co^2+^ titrations

All operations were carried out under anaerobic conditions of a glovebox. Purified, lypoholized Rev was denatured and reduced in 10 mM Tris-HCl, 8 M urea, 2 mM DTT, 1mM EDTA, pH7.5 and refolded on a SP-Sepharose column. The refolded Rev was eluted with 10 mM Tris-HCl, 3M NaCl, pH 7.5. Samples for thiol determination were prepared anaerobically and quantitated upon removal from the glovebox. 0.8 mL of 180 mM reduced apo-Rev was loaded into an anaerobic cuvette. About 140 mL of 0.91 mM Co^2+^ was taken up in an adjustable volume Hamilton syringe and attached to the cuvette via seals and brought out of the glovebox. Optical spectra of the apo-protein and following each addition of a small aliquot (10-15 mL) of Co^2+^ solution were collected on a Hewlett-Packard 8452A spectrometer. The spectra were corrected for light scatter.

#### Fitting

The titration was fit with an equation that accounts for the depletion of both free components, fitting both K_d_ and the total protein concentration (see Materials and Methods in the Supplementary Material). The resulting fit gave an apparent K_d_ of 0.3 uM and a total protein concentration of 13 uM. The P_tot_ value falls between the monomer and dimmer concentration, suggesting that the Rev dimerization is intermediate at the 17.3 uM monomer concentration used in these studies, and Zn^2+^ binding must shift that equilibrium. Further, there is a systematic deviation of the data points below the curve al low Zn^2+^ concentrations and above the curve at higher Zn^2+^ concentrations just below the saturation asymptote, indicative of cooperative binding. We attribute this effect to the promotion of dimers that enter Rev filaments via an isodesmic model. At low Zn^2+^ concentrations (<10uM) stoichiometric complexes must form, given the high concentration of protein with respect to the K_d_. Extrapolation of the curve to full saturation indicates that ∼0.7 Zn^2+^ ions bind per Rev monomer, consistent with binding to a Rev dimer.

**Figure S1.**
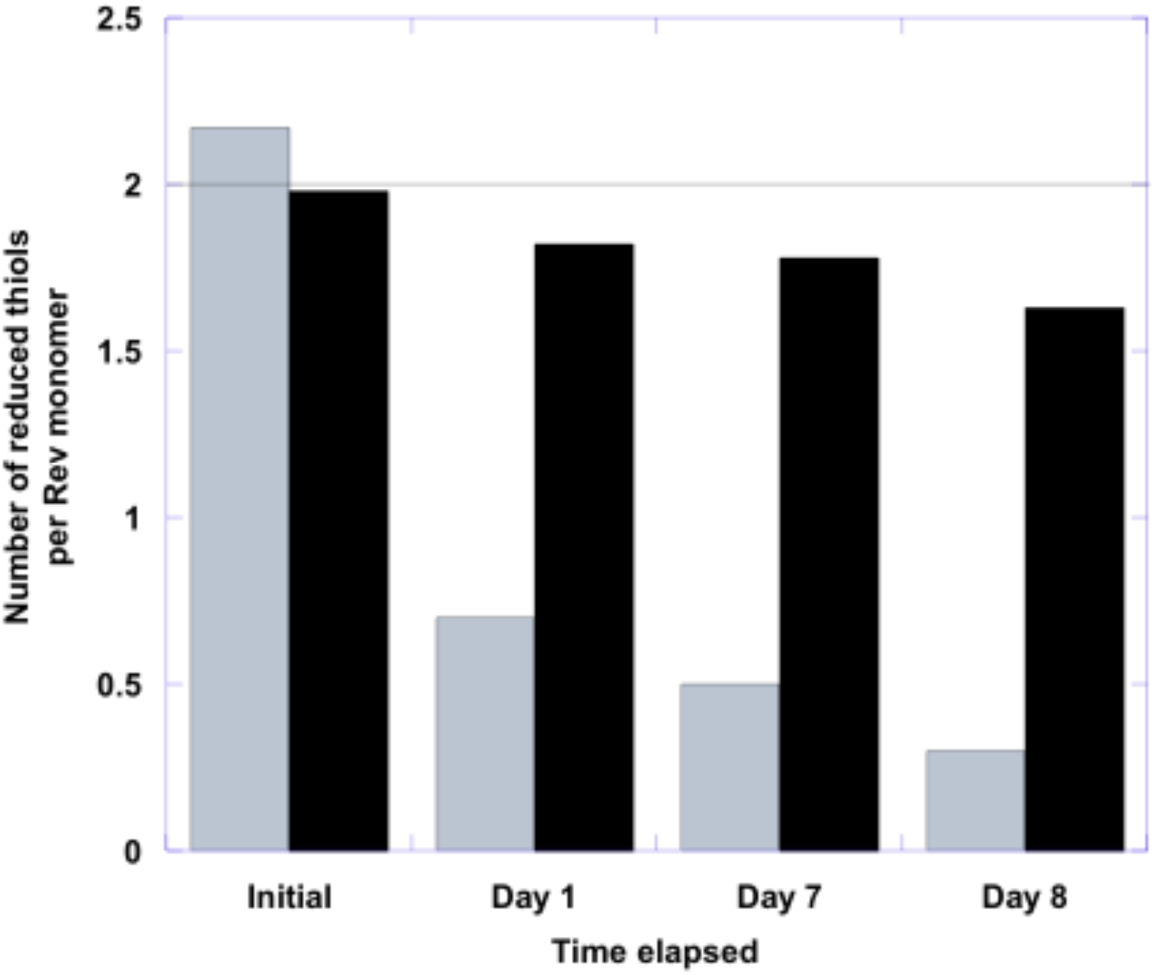
Presence of Zn^2+^ protects Rev thiols from air oxidation *in vitro*. The black bar represents the Zn^2+^Cys_4_ Rev dimer and the gray bars represents apo-Rev.

**Table S1.** Results of Zn^2+^ quantitation by Atomic Absorption Spectroscopy.

| **Protein** | **Zn<sup>2+</sup> addition** | **Final Dialyzate** | **Zn<sup>2+</sup> equivalents** |
| --- | --- | --- | --- |
| Rev* | after refolding | Buffer A** | 0.51 +/- 0.05 |
| RevM10* | after refolding | Buffer A | 0.56 +/-0.06 |
| Rev+ | during refolding | Buffer A | 0.61 +/-0.06 |
| Rev++ | during refolding | Buffer A+EDTA | 0 |

\*\*Buffer A=10 mM Tris-HCl, 1 M NaCl, pH 7.5.
\*these protein samples were refolded in the absence of Zn<sup>2+</sup> and stored in the presence of 2 mM DTT. One equivalent Zn<sup>2+</sup> was added, followed by exhaustive dialysis into Buffer A.
+these protein samples were refolded in the presence of Zn<sup>2+</sup> and 2 mM DTT, followed by exhaustive dialysis into Buffer A.
++these protein samples were refolded in the presence of Zn<sup>2+</sup> and 2 mM DTT, followed by exhaustive dialysis into Buffer A + 2 mM DTT, 1 mM EDTA, pH 7.5.
Values are the average of at least 2 measurements at two protein concentrations of 5 uM and 3 uM .

## Notes

### Competing Interest Statement

The authors have declared no competing interest.

